# Prevalence of pulmonary tuberculosis among patients presenting with cough of any duration in Addis Ababa, Ethiopia

**DOI:** 10.1101/622464

**Authors:** Aynye Negesse, Mulugeta Belay, Girmay Medhin, Sosina Ayalew, Adane Mihret, Mengistu Legesse

**Affiliations:** Federal Ministry of Health, Addis Ababa, Ethiopia; Institute of Health and Society, University of Oslo; Aklilu Lemma Institute of Pathobiology, Addis Ababa University, Addis Ababa, Ethiopia; Armauer Hansen Research Institute, Addis Ababa, Ethiopia

**Keywords:** Cough less than 2 weeks, Prevalence, PTB, RD9 deletion typing, Ethiopia

## Abstract

**Background:** The current practice in Ethiopia to diagnose tuberculosis is screening patients with cough for at least two weeks. A health facility based study was conducted to estimate the prevalence of smear and culture positive pulmonary TB among patients presenting with cough ≥2 weeks and <2 weeks in Addis Ababa, Ethiopia.

**Methods:** A cross-sectional study design was used to recruit patients with cough of any duration from four selected health centers in Addis Ababa, between August and December 2016. Sputum samples were collected from patients reporting productive cough of any duration and screened for Pulmonary Tuberculosis (PTB) using smear microscopy and culture methods. *Mycobacterium tuberculosis* isolates obtained from culture positive samples were characterized using RD9 deletion typing.

**Results:** Majority (39.7%) of the 725 study participants was in the age range of 20-30 years, and 5.0% were smear positive using smear microscopy. The prevalence of smear positive PTB among patients presented with cough duration of ≥2 weeks was significantly higher compared to those patients presented with cough duration of <2 weeks (10.9% versus 0.7%; χ2=38.98; p=0.001). Using culture method, a total of 86 (11.9%) participants were positive for mycobacteria, and the prevalence (14.6%) of PTB among patients presented with cough duration of ≥2 weeks was not significantly higher compared to prevalence (9.9%) in those patients presented with cough duration of <2 weeks (χ2=3.63; p=0.057). Molecular characterization of 86 culture positive mycobacterial isolates showed that 41 were infected with *Mtb;* 19(46.3%) from those who had cough duration of <2 weeks and 22(53.7%) from those who had cough duration of ≥2 weeks.

**Conclusion:** Screening of PTB using smear microscopy alone and cough duration of at least two weeks would negatively affect early diagnosis and treatment initiation in a considerable number of PTB patients who reports cough duration of <2 weeks with the potential of contributing to the spread of TB. Therefore, screening of patients with cough of any duration using both smear microscopy and culture methods is likely to contribute to the success of any effort towards the control of TB.

## Background

Tuberculosis (TB) is responsible for ill health among millions of people each year in low income countries [1]. An estimated 10.4 million new TB cases, 1.3 million TB deaths and an additional 0.37 million deaths as a result of TB disease among HIV-infected people were reported globally in 2016 [1]. Most of the estimated number of incident cases in 2016 occurred in the WHO South-East Asia Region (45%), the WHO African Region (25%) and the WHO Western Pacific Region (17%) [1]. However, TB mortality rate is falling at about 3% per year and TB incidence is falling at about 2% per year [2].

In Ethiopia various efforts have been made to control TB since 1950s, when TB was recognized as a major public health problem [3]. Despite all these national and international efforts, TB still remains one of the major public health problems in the country [1].

The recommended strategy to control TB in low income countries including Ethiopia, where 95% of the TB cases occur, is to detect and promptly treat smear-positive cases [4]. It is known that delayed diagnosis results in more extensive disease, more complications and leads to a higher mortality [5]. It also leads to an increased period of infectivity in the community [6].

To prevent further spread of infection among families and communities, early detection of PTB cases is very important in the TB control [6]. Currently, screening for TB in Ethiopia is limited to patients presenting with cough lasting at least for 2 weeks. On the other hand, a long delay in the diagnosis of TB is a well-documented problem in the management of TB [7–9]. Previous studies also indicated that excluding patients who reported cough lasting less than 2 weeks from screening for TB leads to continued transmission to others and delayed diagnosis [10, 11]. Therefore, this study was conducted to provide additional evidence on the prevalence of PTB among patients presented with cough duration of <2 weeks and among patients presented with cough duration of ≥ 2 weeks using smear microscopy and culture methods. Study participants were recruited among patients attending four health centers of Addis Ababa, Ethiopia.

## Methods

### Study area and participants

This cross sectional survey was conducted from August to December 2016 in four selected health centers of Akaki kality and Nefas silk Lafto Sub-cities in Addis Ababa. Addis Ababa is the capital and chartered city of Ethiopia. Administratively, Addis Ababa has 10 sub-cities which are further divided into 116 districts. Akaki Kality and Nifas Silk-Lafto are two of the 10 sub-cities of Addis Ababa in which the current study health centers are located.

According to the 2017 population projection of Ethiopia, the total population of Akaki Kality subcity is 227,182 and that of Nefassilk Lafto sub-city is 396,486 [12]. The study participants consisted of individuals whose age is above 12 years presented to the four health centers and considered to be part of the study.

### Sample size estimation

The sample size was estimated using the following assumptions: a 21.3% prevalence of smear positive PTB among patients who will present with cough duration of ≥2 weeks [13] and 1.9% prevalence of smear positive PTB among patients who will present with cough duration of <2 weeks [11], 4% margin of error and 95% confidence level. With these assumptions, 302 individuals with cough lasting for ≥2 weeks and 423 individuals with cough lasting <2 two weeks were required to estimate the prevalence in each of the two groups.

### Collection of questionnaire based information and sputum samples

A questionnaire was used to collect relevant data from PTB suspects recruited from the outpatient departments of the four health centers and consented to be part of the study. Health workers identified from each study sites were given responsibility of registering patients whose age is at least 12 years with productive cough of any duration. Consenting the identified potential study participant and requesting the consenting study participants to submit two sputum samples (spot-spot schedule) as per the national guidelines [14]. Socio-demographic characteristics of the study participants including age, sex, marital status, occupation, education, duration of cough, history of TB, history of close contact with TB patients, history of khat chewing, alcohol consumption and cigarette smoking were documented on the questionnaire. PTB suspects were also interviewed about various symptoms such as fever, chest pain, weight loss and loss of appetite.

### Processing, transportation and laboratory analysis of sputum sample

A portion of the sputum samples collected from each individual was processed using Ziel-Neelsen staining (ZN) technique for smear microscopy and examined on the same day to identify acid-fast bacilli (AFB) [15]. The remaining portion of sputum sample was transported to TB laboratory of Aklilu Lemma Institute of Pathobiology (ALIPB) using a cold box and stored at −80°C until processed for culture as previously described [16].

About 0.5 ml of neutralized sputa was inoculated onto test tube containing Lowenstein-Jensen culture medium with 12.5ml glycerol or 0.4% w/v pyruvate and incubated for 8 weeks. Cultures were considered negative when no colonies were seen after 8 weeks incubation period [17]. In case of culture with visible colonies [18], ZN staining was performed to confirm the presence of AFB using microscopy. AFB positive isolates were heat-killed at 80°C for 1 hour using water bath, and stored at −20°C until molecular characterization was performed. *Mtb* isolates were identified using PCR-based genotyping for RD9 deletion as previously described [19].

The PCR amplification mixtures used for RD9 deletions were as follows: reactions were performed in a total volume of 20 μl consisting of 10 μl HotStarTaq Master Mix (Qiagen, United Kingdom), 7.1 μl distilled H2O, 0.3 μl of each oligonucleotide primer (100 μM), and 2 μl DNA template. The Oligonucleotide primers used were: RD9 Flank F: AACACGGTCACGTTGTCGTG; RD9 Flank R: AACACGGTCACGTTGTCGTG and RD9 Internal R: TTGCTTCCCCGGTTCGTCTG. The mixture was heated in a thermal Cycler (Gene AMP 9700) for 15 minutes at 95°C and then subjected to 35 cycles of one-minute duration at 95°C, one minute at 55°C, one minute at 72°C and 10 minutes at 72°C. A molecular weight of 396bp was considered as *Mtb*, while a molecular weight 575bp was considered as *M. bovis* [20].

For gel electrophoresis, 8 μl PCR products was mixed with 2 μl loading dye, loaded onto 1.5% agarose gel and electophoresed at 100 V and 500 mA for 45 min. The gel was then visualized using a computerized Multi-Image Light Cabinet (VWR). *Mtb* H37Rv, *M. bovis* bacille Calmette-Guérin, and water were included as positive and negative controls. Interpretation of the result was based on bands of different sizes as a molecular weight of 396bp was considered as *Mtb*, while a molecular weight 575bp was considered as M. bovis.

Gel electrophoresis was used for the separation of the PCR product. For gel electrophoresis, 10 μl PCR product was mixed with 2 μl loading dye, loaded onto 3% agarose gel and electrophoresed at 110 V and 400 mA for 30 min. The gel was then visualized using a computerized Multi-Image Light Cabinet (VWR). *Mtb* H37Rv, *M. bovis* bacille Calmette-Guérin, and water were included as positive and negative controls. Interpretation of the result was based on bands of different sizes, as previously described [20].

### Data management and processing

Data was computerized using Epi Data version 3.1 and analysis was performed using Stata version 12. The proportion of patients with smear positive PTB was calculated for the two study groups (i.e. according to their cough duration). A possible association between positivity for PTB and patients’ background characteristics as well as duration of cough was investigated using bivariate and multivariable logistic regression. Statistically significant association was reported whenever p-value was less than 5%.

## Results

### Associations of sociodemographic characteristics of the study participants with prevalence of PTB confirmed by smear and culture methods

Socio-demographic characteristics of 725 study participants and their associations with prevalence of PTB are summarized in Table 1. The oldest study participant was 90 years, the mean age was 36.7 years and 39.7% were in the age range of 20-30 years.

**Table 1:**
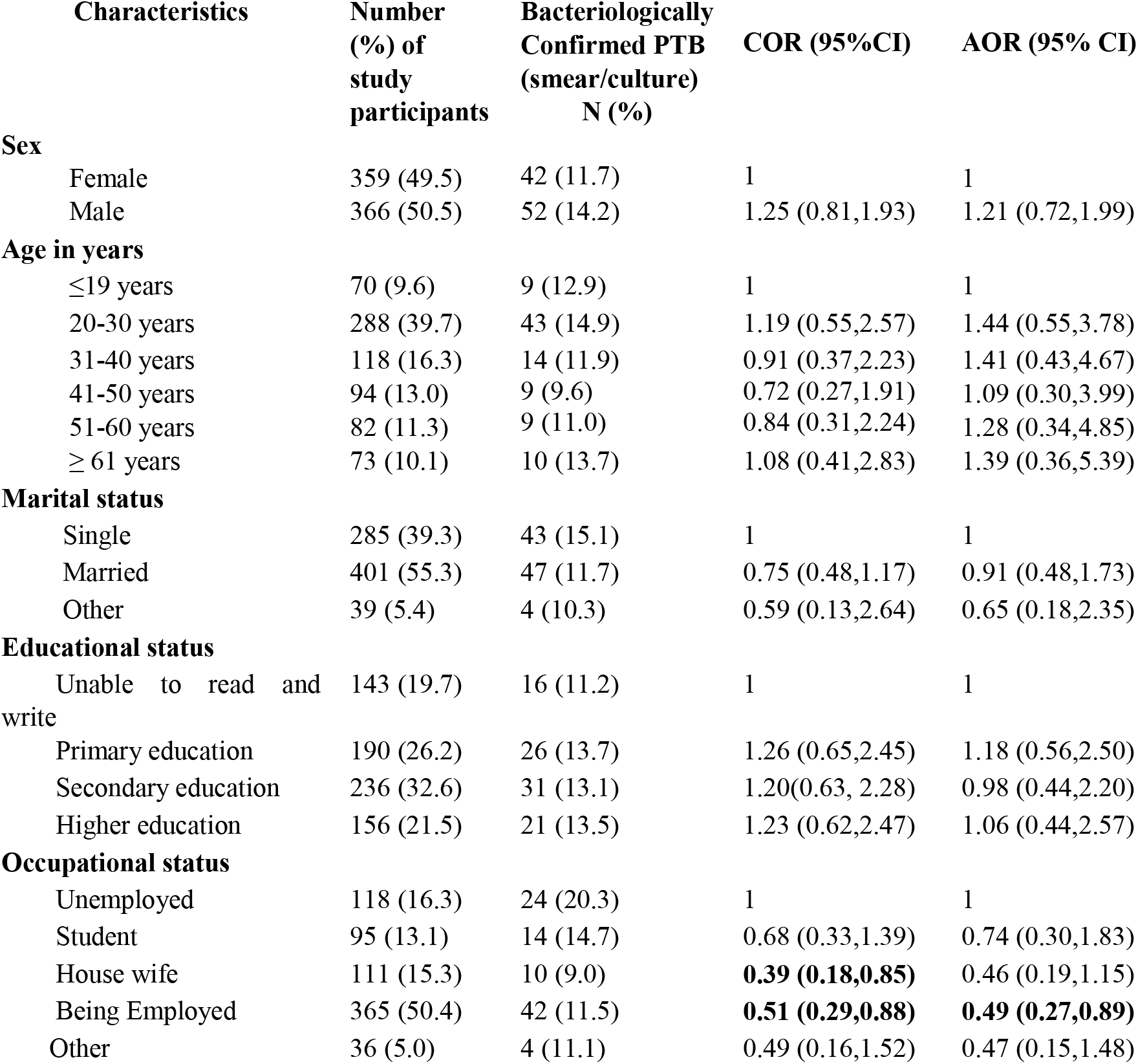
Background characteristics of the study participants and prevalence of PTB

| Characteristics | Number (%) of study participants | Bacteriologically Confirmed PTB (smear/culture) N (%) | COR (95%CI) | AOR (95% CI) |
| --- | --- | --- | --- | --- |
| **Sex** |  |  |  |  |
| Female | 359 (49.5) | 42 (11.7) | 1 | 1 |
| Male | 366 (50.5) | 52 (14.2) | 1.25 (0.81,1.93) | 1.21 (0.72,1.99) |
| **Age in years** |  |  |  |  |
| $\leq 19$ years | 70 (9.6) | 9 (12.9) | 1 | 1 |
| 20-30 years | 288 (39.7) | 43 (14.9) | 1.19 (0.55,2.57) | 1.44 (0.55,3.78) |
| 31-40 years | 118 (16.3) | 14 (11.9) | 0.91 (0.37,2.23) | 1.41 (0.43,4.67) |
| 41-50 years | 94 (13.0) | 9 (9.6) | 0.72 (0.27,1.91) | 1.09 (0.30,3.99) |
| 51-60 years | 82 (11.3) | 9 (11.0) | 0.84 (0.31,2.24) | 1.28 (0.34,4.85) |
| $\geq 61$ years | 73 (10.1) | 10 (13.7) | 1.08 (0.41,2.83) | 1.39 (0.36,5.39) |
| **Marital status** |  |  |  |  |
| Single | 285 (39.3) | 43 (15.1) | 1 | 1 |
| Married | 401 (55.3) | 47 (11.7) | 0.75 (0.48,1.17) | 0.91 (0.48,1.73) |
| Other | 39 (5.4) | 4 (10.3) | 0.59 (0.13,2.64) | 0.65 (0.18,2.35) |
| **Educational status** |  |  |  |  |
| Unable to read and write | 143 (19.7) | 16 (11.2) | 1 | 1 |
| Primary education | 190 (26.2) | 26 (13.7) | 1.26 (0.65,2.45) | 1.18 (0.56,2.50) |
| Secondary education | 236 (32.6) | 31 (13.1) | 1.20(0.63, 2.28) | 0.98 (0.44,2.20) |
| Higher education | 156 (21.5) | 21 (13.5) | 1.23 (0.62,2.47) | 1.06 (0.44,2.57) |
| **Occupational status** |  |  |  |  |
| Unemployed | 118 (16.3) | 24 (20.3) | 1 | 1 |
| Student | 95 (13.1) | 14 (14.7) | 0.68 (0.33,1.39) | 0.74 (0.30,1.83) |
| House wife | 111 (15.3) | 10 (9.0) | **0.39 (0.18,0.85)** | 0.46 (0.19,1.15) |
| Being Employed | 365 (50.4) | 42 (11.5) | **0.51 (0.29,0.88)** | **0.49 (0.27,0.89)** |
| Other | 36 (5.0) | 4 (11.1) | 0.49 (0.16,1.52) | 0.47 (0.15,1.48) |

Based on the smear microscopy examination, the overall prevalence of smear-positive PTB was 5.0%. Thirty three (10.9%) (95% CI: 7.4 – 14.5) smear positive PTB cases were among those who reported cough duration of ≥2 weeks, and 3 (0.7%) (95% CI: −0.09 – 1.5) smear positive PTB cases were among those who had cough duration of <2 weeks. This resulted in 49/100,000 prevalence of smear positive PTB cases among health seeking population.

A total of 86 (11. 9%) were identified as PTB patients based on the culture result with a contamination rate of 3.8%. Out of these, 28 were among smear positives and 58 were among smear negatives. Additional eight suspects who were reported as smear positive were turned out to be culture negative. Among those who reported cough duration of ≥2 weeks, 44 (14.6%) (95% CI: 10.6 – 18.6) were culture positive. Whereas, 42 (9.9%) (95% CI: 7.1 – 12.8) were among those who had cough duration of <2 weeks. Hence, the prevalence of PTB among health seeking population using culture method was 117/100,000.

After adjusting for potential confounding variables, being employed (AOR = 0.49; 95% CI, 0.27, 0.89) was significantly associated with low PTB positivity.

### Clinical features, other chronic diseases and addictive habits reported by the study participants

Majority (80.1%) of the study participants complained of having more than one symptom suggestive of PTB in addition to cough including fever (65.7%), chest pain (61.2%), night sweats (61.1%) and loss of appetite (59.6%). The reported duration of cough varies between 4 and 365 days, with a median of 10 days. More than half of the study participants (58.3%) reported short duration of cough with a mean duration of 16.6 days (SD ± 20.7) (Table 2).

**Table 2:** Clinical features, other chronic diseases and addictive habits of the study participants and prevalence of PTB

| Characteristics | Number (%)<br>of study<br>participants | Bacteriologically<br>confirmed PTB<br>(smear/culture)<br>N (%) | COR (95%CI) | AOR (95% CI) |
| --- | --- | --- | --- | --- |
| **Fever** |  |  |  |  |
| **No** | 249 (34.3) | 28 (11.2) | 1 | 1 |
| **Yes** | 476 (65.7) | 66 (13.9) | 1.27(0.79,2.04) | 1.31 (0.89,2.16) |
| **Chest pain** |  |  |  |  |
| **No** | 281 (38.8) | 34 (12.1) | 1 | 1 |
| **Yes** | 444 (61.2) | 60 (13.5) | 1.14 (0.72,1.78) | 0.90 (0.55, 1.45) |
| **Night sweating** |  |  |  |  |
| No | 282 (38.9) | 25 (8.9) | 1 | 1 |
| Yes | 443 (61.1) | 69 (15.6) | **1.90 (1.17,3.08)** | **1.71 (1.02, 2.85)** |
| **Loss of appetite** |  |  |  |  |
| No | 293 (40.4) | 38 (13.0) | 1 | 1 |
| Yes | 432 (59.6) | 56 (13.0) | 0.99 (0.64,1.55) | 0.88 (0.55, 1.39) |
| **Duration of cough** |  |  |  |  |
| < 2 weeks | 423 (58.3) | 43 (10.2) | **1** | 1 |
| ≥ 2 weeks | 302 (41.7) | 51 (16.9) | **1.79 (1.16,2.78)** | **1.69 (1.08, 2.66)** |
| **Close contact with known TB patients** |  |  |  |  |
| No | 669 (92.3) | 85 (12.7) | 1 | 1 |
| Yes | 56 (7.7) | 9 (16.1) | 1.32 (0.62,2.78) | 1.33 (0.62,2.85) |
| **Previous history of TB** |  |  |  |  |
| No | 672 (93.0) | 83 (12.4) | 1 | 1 |
| Yes | 51 (7.0) | 11 (21.6) | 1.95 (0.96,3.95) | 1.77 (0.85,3.66) |
| **History of chronic diseases** |  |  |  |  |
| No | 671 (92.5) | 87 (13.0) | 1 | 1 |
| Yes | 54 (7.5) | 7 (13.0) | 1.00 (0.44,2.28) | 0.84 (0.36, 1.96) |
| **History of smoking** |  |  |  |  |
| No | 691 (95.3) | 88 (12.7) | 1 | 1 |
| Yes | 34 (4.7) | 6 (17.7) | 1.47 (0.59,3.65) | 0.64 (0.16,2.54) |
| **History of khat chewing** |  |  |  |  |
| No | 693 (95.6) | 87 (12.6) | 1 | 1 |
| Yes | 32 (4.4) | 7 (21.9) | 1.95 (0.82,4.64) | 2.03 (0.55,7.52) |
| **History of alcohol consumption** |  |  |  |  |
| No | 657 (90.6) | 81 (12.3) | 1 | 1 |
| Yes | 68 (9.4) | 13 (19.1) | 1.68 (0.88,3.21) | 1.42 (0.63,3.19) |

Fifty-six respondents (7.7%) reported that they had close contact with known TB patients in the last 15 years (1.8%) and for a minimum of 15 days (1.8%). Previous history of TB and history of chronic diseases were reported by 51 (7.1%) and 54 (7.5%) respondents, respectively (Table 2).

Based on bacteriologic results (smear microscopy and/or culture), a total of 94 (13.0%) were diagnosed as PTB cases resulting in a prevalence of 128/100,000 among health seeking population. Among patients who coughed for a duration of ≥2 weeks, 51 (16.9%) (95% CI: 12.6% to 21.1%) had bacteriologically confirmed PTB. Similarly, from those who coughed for a duration of <2 weeks, 43 (10.2%) (95% CI: 7.3 – 13.1) had bacteriologically confirmed PTB.

Duration of cough for ≥2 weeks (AOR = 1.69; 95% CI, 1.08-2.66) and night sweating (AOR = 1.71; 95% CI, 1.02 – 2.85) were significantly associated with bacteriologically confirmed PTB positivity (Table 2).

### Summary results from RD9 deletion typing

Molecular characterization of 86 mycobacterial isolates using RD9 deletion typing showed that 41 (47.7%) isolates contained an intact RD9 implying that the 41 isolates belonged to *Mtb*. Among these 41 patients, 19 (46.3%) reported cough duration of <2 weeks and 22 (53.7%) reported cough duration of ≥2 weeks. Among the 28 smear positive PTB cases, 18 (64.3%) were confirmed as *Mtb* by RD9 deletion typing. (Table 3)

**Table 3:** Associations of variables with RD9 deletion typing results

| Characteristics | Number<br>(%) of<br>study<br>participants | RD9 deletion<br>typing<br>positive N (%) | COR (95%CI) | AOR (95%CI) |
| --- | --- | --- | --- | --- |
| **Sex** |  |  |  |  |
| Female | 37 (43.0) | 19 (51.3) | 1.00 | 1.00 |
| Male | 49 (57.0) | 22 (44.9) | 0.77 (0.55, 1.82) | 1.27 (0.41, 3.92) |
| **Age in years** |  |  |  |  |
| ≤ 19 years | 7 (8.1) | 1 (14.3) | 1.00 | 1.00 |
| 20-30 years | 39 (45.4) | 22 (56.4) | 7.76 (0.85, 70.75) | 5.52 (0.52, 58.16) |
| 31-40 years | 13 (15.1) | 13 (53.8) | 7.00 (0.65, 75.73) | 6.82 (0.42, 110.77) |
| 41-50 years | 9 (10.5) | 5 (55.6) | 7.50 (0.62, 90.65) | 8.44 (0.37, 191.01) |
| 51-60 years | 8 (9.3) | 4 (50.0) | 6.00 (0.48, 75.34) | 9.55 (0.29, 317.26) |
| ≥ 61 years | 10 (11.6) | 2 (20.0)) | 1.50 (0.11, 20.68) | 1.77 (0.06, 52.46) |
| **Marital status** |  |  |  |  |
| Never married | 38 (44.2) | 18 (47.4) | 1.00 | 1.00 |
| Ever Married | 48 (55.8) | 23 (47.9) | 1.02 (0.44, 2.40) | 1.38 (0.30, 6.24) |
| **Educational status** |  |  |  |  |
| No formal education | 16 (18.6) | 6 (37.5) | 1.00 | 1.00 |
| Formal education | 70 (81.4) | 35 (50.0) | 1.67 (0.55, 5.08) | 1.93 (0.32, 11.63) |
| **Duration of cough** |  |  |  |  |
| < 2 weeks | 42 (48.8) | 19 (45.2) | 1.00 | 1.00 |
| ≥ 2 weeks | 44 (51.2) | 22 (50.0) | 1.21 (0.52, 2.83) | 1.55 (0.54, 4.40) |
| **Close contact with known TB patients** |  |  |  |  |
| No | 77 (89.5) | 38 (49.3) | 1.00 | 1.00 |
| Yes | 9 (10.5) | 3 (33.3) | 0.51 (0.12, 2.20) | 0.56 (0.10, 3.10) |
| **Previous history of TB** |  |  |  |  |
| No | 76 (88.4) | 35 (46.0) | 1.00 | 1.00 |
| Yes | 10 (11.6) | 6 (60.0) | 1.76 (0.46, 6.73) | 1.57 (0.32, 7.69) |
| **History of chronic diseases** |  |  |  |  |
| No | 80 (93.0) | 39 (48.7) | 1.00 | 1.00 |
| Yes | 6 (7.0) | 2 (33.3) | 0.53 (0.09, 3.03) | 0.44 (0.06, 3.18) |

## Discussion

In this study, a health facility based prevalence of PTB is reported using smear microscopy and culture methods among patients presenting to the facility with cough of any duration. Based on the smear microscopy examination the overall prevalence of smear-positive PTB was 5.0%, and the prevalence was higher among patients who reported cough for ≥2 weeks (10.9%) compared to those patients who reported cough for <2 weeks (0.7%). The smaller number of positive cases detected among patients with cough less than two weeks should not be considered insignificant since these individuals would contribute to the transmission of the disease among the families and the community if they were not screened and treated [9, 21]. According to the manual of Tuberculosis, Leprosy and TB/HIV Prevention and Control Program in Ethiopia, any person with cough duration of two weeks or more is usually suspected for PTB and screened as per the guideline [14]. This practice excludes patients who complained cough for less than two weeks causing delay in the diagnosis of potential cases thereby increasing the risk of transmission to others. Previous studies also identified smear positive PTB cases among patients who reported cough for duration of <2 weeks [10, 11, 22].

Based on culture result, the overall prevalence of PTB was 11.9% which varies depending on reported duration of cough. The observed prevalence among patients with cough for <2 weeks was comparable with patients who had cough for ≥2 weeks. Majority of culture positives were from smear negative cases which is similar to the findings observed in Malawi among patients with short duration of cough [22].

Bacteriologically confirmed PTB among study participants with short duration was 10.2% which is in line with a study in China in which acute cough was independent predictors of bacteriologically positive TB diagnosis [23]. The prevalence in the current study is higher than the previous report in Eastern Ethiopia [24] and elsewhere [22]. However, the prevalence of bacteriologically confirmed PTB in the current study among patients with longer duration of cough is lower than the previous findings from different parts of Ethiopia [11, 21, 25] and elsewhere [22]. The observed difference might be the variation of the study participants in which this study screened all patients who had cough of any duration in our study.

The findings of the current study supports the previous findings that showed suspects who were reported as smear positive turned out to be culture negative [21, 22, 26, 27], this might be partly be explained by reduced recovery of *Mtb* from cultures result based on specimen stored a very long time [28,29] or due to a rather harsh decontamination process [29], or could be false smear positive report due to technical problems.

In the current study duration of cough and night sweating were identified as potential risk factors for being PTB positive and currently employed was identified as a significant protective factor. In previous study unemployment was independently and significantly associated with being TB positives [23]. Employed individuals perhaps can get information from co-workers which helps them to make informed decision and visit health institutions on a timely manner when they feel having symptoms and diagnosed TB. In Tanzania [10] and in Malawi [28] smear positive PTB was similar among study participants who had coughed for less than two weeks and among those who had cough for at least two weeks.

The proportion of culture positive samples that gave positive signal to RD9 deletion typing in the study is significantly lower than the proportion reported in the molecular characterization of *Mycobacterium tuberculosis* complex study in Gambella Regional state of Ethiopia [32] and elsewhere [39].

Not being able to further characterize 45 isolates for *nontuberculous mycobacterial* (NTM) infection because of shortage of reagents is one of the limitations of the current study. The second limitation is potential recall bias linked to duration of cough since patients remember quite often the recent most cough duration. Culture was not done for each sample within three days as recommended by World Health Organization and this could potentially affect the prevalence of culture positivity.

## Conclusion

The results of this study indicate that screening of PTB using smear microscopy alone and use of cough duration for at least two weeks have a potential negative effect on early diagnosis and timely treatment initiation in a considerable number of PTB patients who reports cough for <2 weeks, and contributes to the spread of TB. Higher proportion of smear negative TB suspects was culture positive which showed that the current diagnostic method (smear microscopy) in the health facilities resulted in delayed in seeking care and in diagnosis thereby increasing the risk of transmission to others. Lower proportion of Mtb was identified by RD9 deletion typing. This suggests the need for further studies to identify the contribution of NTM for TB.

Therefore, screening of patients with cough of any duration using both smear microscopy and culture methods is very important for the control of TB. For the current case finding strategy in Ethiopia, the combined CXR and symptom screening in addition to cough should be done helping to improve the screening yield in different procedures. Furthermore, there is a need for improving TB lab at each health care level such as new diagnostic tools including LED, culture and immunological test.

## List of abbreviations

AHRI: : Armauer Hansen Research Institute
ALIPB: : Aklilu Lemma Institute of Pathobiology
AOR: : Adjusted Odds Ratio
CFR: : Case Fatality Rate
CI: : Confidence Interval
COR: : Crude Odds Ratio
CSA: : Central Statistics Agency
DNA: : Deoxyribonucleic Acid
FMOH: : Federal Ministry of Health
HBC: : High Burden Country
HIV+TB: : Human Immunodeficiency Virus-tuberculosis co-infection
LJ: : Lowenstein-Jensen
MtbC: : *Mycobacterium tuberculosis* complex
NSL: : Nefas Silik Lafto Sub-city
PCR: : Polymerase chain reaction
PTB: : Pulmonary tuberculosis
RD9: : Region of difference 9
RNAse: : Ribonucleic Acidase enzyme
SD: : Standard deviation
TB: : Tuberculosis

## Declarations

### Ethics approval and consent to participate

The study protocol was approved by the Institutional Review Board of ALIPB, Addis Ababa University and Ethics Committee of Addis Ababa Health Bureau before starting data collection. Written informed consent was obtained from each participant (guardians for children) after describing the objective of the study to each participant/guardian before enrolment into the study. As part of the routine clinical care, all patients with smear positive PTB were referred to the TB clinic for treatment and smear negative patients with suggestive diagnosis of TB were treated according to the national guideline. Furthermore, the health centers physicians/health officers/nurses were informed about those patients who were found positive by the culture method since OPD health workers have the mandate of referring TB patients to TB clinic for appropriate management and patients were contacted.

### Consent to publication

Consent was obtained from each participant after describing the objective of the study to publication.

### Availability of data and materials

The datasets used and/or analyzed during the current study are available from the corresponding author on reasonable request.

### Competing interests

The authors declare that they have no competing interests.

### Funding

I declared that Addis Ababa University supported me in the process of designing of the study and data collection.

### Authors’ contributions

This paper was based on the Masters thesis of AN; he collected the data, analyzed and interpreted the result from sputum specimen. ML, MB, GM and AM were the supervisors in the whole process of the thesis, and were a major contributor in writing the manuscript. SA was contributing in assisting the molecular procedures. All authors read and approved the final manuscript.

## Acknowledgement

We raise our deepest gratitude to Aklilu Lemma Institute of Pathobiology, Addis Ababa University. We are extending our appreciation to Addis Ababa Health Bureau and Akaki Kality Health Office for providing permission to do this research in the selected health facilities. We are grateful to outpatient department case team coordinators and laboratory technicians at the study health facilities for their sincere efforts in collecting the data. We are indebted to the study participants for their willingness and cooperation during the study. Aklilu Lemma Pathobiology TB Laboratory (Professor Gobena Ameni) and Armauer Hansen’s Research Institute are highly appreciated for providing permission to do culture and molecular procedures; laboratory professionals particularly Mr Hailu Getu who were in our side every time, Samuel Tolosa and Aboma Zewude gave us unpayable and consistent support.

## Author’s information

1 Aklilu Lemma Institute of Pathobiology, Addis Ababa University, P.O. Box 1176, Addis Ababa, Ethiopia.

2 Institute of Health and Society, University of Oslo

3 Armauer Hansen Research Institute, Addis Ababa, Ethiopia

